# Quorum sensing antiactivators constrain *Pseudomonas aeruginosa* RhlR activity

**DOI:** 10.1101/2025.10.31.685934

**Authors:** Varun Sridhar, Nicole. E. Smalley, Ajai A. Dandekar, Kyle L. Asfahl

**Affiliations:** Department of Plant and Microbial Biology, University of California - Berkeley, Berkeley, CA 94720; Department of Microbiology, University of Washington, Seattle, WA 98195; Department of Medicine, University of Washington, Seattle, WA 98195; Microbial Interactions and Microbiome Center, University of Washington, Seattle, WA 98195

## Abstract

*Pseudomonas aeruginosa*, an opportunistic pathogen, uses a cell-cell communication system called quorum sensing (QS) to regulate gene expression in response to population density. *P. aeruginosa* QS involves, in part, two transcription factors, LasR and RhlR, that respond to *N*-acyl homoserine lactone (AHL) signals. Two proteins known as “antiactivators,” QteE and QslA, attenuate QS by inhibiting LasR, RhlR, or both. While initial characterization of antiactivation has revealed the considerable influence these factors may have on dampening QS, details regarding the individual impacts of these antiactivators on *P. aeruginosa* QS activity remain scant. Additionally, the effects of antiactivators on RhlR QS activity and in QS systems in isolates or strain lacking LasR have yet to be explored. To investigate how QteE and QslA each modulate LasR or RhlR independently, we combined gene deletion and over-expression analysis of each antiactivator in wild-type *P. aeruginosa* (PAO1) and two strains with *rhl*-dominated QS: clinical isolate E90 and PAO Δ*lasR* Δ*mexT*. As measured with a transcriptional reporter, over-expression of *qteE* or *qslA* notably reduced RhlR activity in PAO1 and PAO Δ*lasR* Δ*mexT*, but only expression of *qteE* had a marked effect on RhlR activity in E90. Expression analysis indicates LasR and RhlR repress QteE transcription, but not QslA. By over-expressing *qslA* in the absence of QteE and vice versa, we demonstrate that QslA activity and corresponding effects on QS phenotypes can be QteE-dependent in some scenarios. Our results reveal a nuanced role for individual antiactivator proteins in affecting the layered *P. aeruginosa* QS circuitry.

**Importance:** Quorum sensing (QS) is a cell signaling mechanism that enables populations of *Pseudomonas aeruginosa* to coordinate group behaviors such as biofilm formation, virulence factor production, and antibiotic tolerance once a critical cell-density threshold is reached. *P. aeruginosa* employs two “antiactivator” proteins that attenuate QS at low cell densities, dampening QS activation. The specific effects of individual antiactivators on the complex and hierarchically-arranged *P. aeruginosa* QS systems remain undefined. Here, we use two strains with rewired QS circuits to independently assess the effects of QS antiactivators on each QS circuit. We find that while one antiactivator selectively targets one QS circuit, the other can broadly target both with strong effects on QS activity. This work reveals an additional layer of complexity to counter-regulation of QS signalling and further defines antiactivation as a mechanism *P. aeruginosa* uses to finely tune QS responses.

## Introduction

Many bacteria use quorum sensing (QS), a signaling system that couples cell-cell communication and gene regulation in response to population density, to concert population-wide behavior (1). A common type of QS in Gram-negative bacteria is mediated by acyl-homoserine (AHL) signal molecules. These diffusible AHL signals are produced by LuxI-type synthases (I-proteins) and received and bound by LuxR-type regulators (R-proteins). In most cases, the signal-bound R-protein serves as a transcriptional activator that binds conserved promoter sequences to control expression of target genes (1, 2).

*Pseudomonas aeruginosa*, an opportunistic pathogen and a model for QS in Gram-negative bacteria, employs two complete LuxI/R-type gene circuits: LasI/R and RhlI/R. The LasI synthase produces an *N*-3-oxo-dodecanoyl-homoserine lactone (3OC12-HSL) signal which binds to the transcription activator LasR, and the RhlI synthase produces *N*-butanoyl-homoserine lactone (C4-HSL) which binds to RhlR (3). At low cell density, basal transcription of I-proteins yields low concentrations of signal that minimally bind and activate R-proteins, yielding an overall QS-off state. At high cell density and signal concentrations, both R-proteins form homodimers upon binding of their cognate signal to produce transcriptionally active complexes and a QS-on state (4, 5). Activation of each QS system is amplified via auto-induction and further transcription of their respective synthases under conditions of high cell density and signal concentration. LasR controls the activation of *rhlR* in lab strains, positioning the two QS systems in a hierarchy with LasR as the master regulator (6). Together, the *las* and *rhl* QS circuits control the transcription of overlapping regulons that include genes encoding a battery of secreted products and virulence factors such as extracellular proteases, rhamnolipids, phenazines, and hydrogen cyanide (7).

A remarkable trait of QS is that a population can trigger the induction of QS circuits in a majority of cells once the group reaches a target cell density, yet avoid early activation in individual cells (1, 2). A natural question therefore arises: how do bacteria prevent early QS activation below a target density threshold? One such mechanism described in several AHL QS systems, including the LasI/R system of *P. aeruginosa*, is inhibition of R-protein activation through binding by so-called “antiactivator” proteins. QS antiactivation was first defined in the AHL QS system of *Agrobacterium tumefaciens*, in which the antiactivator TraM directly binds the receptor-regulator TraR to sequester it and prevent further dimerization and activation of the QS network (8–10). Mutants of TraM display higher QS activity and earlier activation of QS-controlled genes, indicating an important role for antiactivation in delaying QS induction in this system (9). *P. aeruginosa* employs proteins that serve to similarly dampen QS activity, two of which fall under the description of R-protein antiactivation: QteE and QslA (11–13). QteE is known to destabilize LasR dimers independently of transcription or translation, suggesting a direct interaction and sequestration of heterodimers or other post-translational mechanism (11). Some evidence has suggested QteE also targets and serves to destabilize RhlR (11). QslA works in a similar fashion as TraM (despite low sequence similarity), directly binding LasR monomers in a 2:1 ratio to prevent dimerization (12, 13). *qslA* is expressed 55.8 times lower in a *P. aeruginosa* model for hypervirulence compared to the WT laboratory strain; this decreased expression drives higher production of many virulence factors and expression *of lasR* and *rhlR*, indicating repression of antiactivation may promote higher virulence in *P. aeruginosa* infections (14). Deleting both antiactivators has a greater effect on both the timing of QS induction and the magnitude of QS-controlled phenotypes than individual deletions of either gene (15). Measuring *lasI* expression in AHL signal-proficient and -deficient cells grown in coculture, Smith and Schuster demonstrated that signal-proficient cells exhibited much stronger *lasI* induction at low cell densities in an antiactivator mutant background compared to the wild-type, suggesting a clear role for QteE and QslA in prevention of early *las* circuit activation (16).

Although previous work has uncovered some aspects of how these two *P. aeruginosa* antiactivators function to affect QS induction, several questions remain. Most work on the mechanisms underlying QteE and QslA activity has focused on LasR and not RhlR, a consequence of LasR regulating the *rhl* QS circuit in many lab strains. Many strains of *P. aeruginosa* isolated from infections or the environment do not have a functional LasR and instead use RhlR to control QS, underscoring both a need for further study of these divergent QS circuits while also presenting a potential resource (17, 18). How antiactivation affects RhlR activity in QS circuits divorced from LasR remains unknown. The repressive effects of QteE and QslA appear distinct for LasR, but further investigation is warranted to better understand their individual roles in signal reception or LasR activity and stability, in addition to exploration of their effects on RhlR. Here, we leverage both a lab-adapted strain and a clinical isolate that showcase *rhl*-dominated QS to demonstrate a clear antiactivation effect of QteE, and to a lesser extent QslA, on RhlR. Using a combination of deletion mutants and over-expression constructs coupled with measurements of transcriptional activity, signal response, protein expression, and QS phenotypes, we uncover considerable nuance in the role of QteE and QslA in dampening QS in *P. aeruginosa*.

## Results

### Bacterial strains for studying LasR-independent QS circuits

While Las-independent Rhl-QS systems are common in natural isolates of *P. aeruginosa,* laboratory strain genetics have made study of this type of QS challenging. The effects of deleting *qteE, qslA*, or both antiactivator-encoding genes in conjunction have been examined in wild-type *P. aeruginosa*, strain PAO1, but such deletions have not been examined in QS circuits free of the LasR-dominated regulatory hierarchy common in this widely-used laboratory strain (11, 13, 15, 16). Analytical strategies used to address this issue with respect to QteE activity include inducible expression of *rhlR* in a Δ*lasR* Δ*rhlR* mutant, or ectopic expression of the pertinent QS and antiactivator proteins in *E. coli* (11). While both approaches have yielded some insight into QteE’s role in dampening *P. aeruginosa* QS, neither have been employed to better understand QslA, and importantly, these previous efforts have not addressed antiactivation of RhlR in a native *P. aeruginosa* regulatory context. One solution is to use clinical isolates from *P. aeruginosa* infections in people with cystic fibrosis (19–21). Many of these clinical isolates carry nonfunctional alleles of *lasR* and have “rewired” the QS circuits to favor RhlR/I as the dominant QS system relative to the PAO1 paradigm (17, 22, 23). Isolate E90 has been characterized as a genetically tractable strain that carries a nonfunctional *lasR* allele and demonstrates Las-independent RhlR activity and QS phenotypes (17, 24–27). Another solution lies in an *in vitro*-evolved lab strain variant that expresses a subset of the RhlR regulon despite the absence of LasR. Kostylev and colleagues proposed that clinical isolates of *P. aeruginosa* may evolve QS circuitry that favors RhlR-based QS by developing mutations in the LysR-type regulator MexT, supported by their observations of reproducible *mexT* mutation in lab conditions that require QS for growth (28). They demonstrated that targeted deletions of just *mexT* and *lasR* in strain PAO1 produces the same QS regulation phenotype as the evolved strain, resulting in genetically tractable *P. aeruginosa* that expresses some RhlR-regulated genes independent of LasR. Therefore, we selected both the clinical isolate E90 and PAO Δ*lasR* Δ*mexT* as complimentary models for investigation of antiactivation in *rhl*-dominated QS.

### QteE, in contrast to QslA, separately modulates LasR and RhlR

Previous work has demonstrated that inactivating mutations in *qteE* or *qslA* in strain PAO1 advances the induction of genes regulated by both LasR and RhlR (15). In this study, we aimed to broaden this understanding to other native QS circuitries through both mutation and over-expression of antiactivator genes. Here, we analyzed clean deletions in *qteE* or *qslA* in PAO1 alongside *qteE* deletions in PAO Δ*lasR*Δ*mexT* and E90. To control over-expression of *qteE* or *qslA*, we integrated an additional copy of either gene under the control of a synthetic arabinose-inducible promoter into a neutral chromosomal site (P*_araBAD_-qteE*, P*_araBAD_-qslA*) (29). As a readout of RhlR activity, we monitored promoter activity of the rhamnosyl transferase gene *rhlA* with a plasmid-borne GFP-transcriptional reporter (P*_rhlA_-gfp*) over time, and we used this bulk fluorescence data to calculate time to half-maximal GFP expression as a quantitative analysis of induction timing. *rhlA* expression has been widely observed as strictly controlled by RhlR, regardless of strain background, so even in the context of active LasR this promoter provides a reliable index of RhlR activation (30, 31).

In PAO1, deletion of *qteE* predictably advances timing and magnitude of P*_rhlA_-gfp*-derived fluorescence owing to the Las-dependent hierarchy in this strain, while *qteE* over-expression abolishes it (Figure 1A). P*_rhlA_-gfp*-derived fluorescence is generally lower in PAO Δ*lasR* Δ*mexT* and E90 (Figure 1C, 1E). Deletion of *qteE* in these Las-independent strains shows little additional fluorescence over their isogenic parents and does not significantly reduce time to half-maximal induction, but over-expression still completely abolishes P*_rhlA_-gfp* activity in these strains (Figure 1C, 1E). On the other hand, deletion of *qslA* in PAO1 shows slightly earlier P*_rhlA_-gfp* activity, but an overall similar magnitude of expression (Figure 1B). Surprisingly, this activity is not completely abrogated upon induction of P*_araBAD_-qslA*, only delayed. While multiple attempts at construction of *qslA* deletions in both PAO Δ*lasR* Δ*mexT* and E90 proved unsuccessful, we were able to induce overexpression of this gene via P*_araBAD_-qslA*. Expression of *qslA* in PAO Δ*lasR*Δ*mexT* reveals an antiactivation effect similar to PAO1, with modest but significant antiactivation of RhlR and P*_rhlA_-gfp* activity (Figure 1D). The effect of *qslA* over-expression is even weaker in E90 and barely discernable from WT E90 (Figure 1F, not significantly different *p* = 0.5262). Together, these data show that in addition to the strong antiactivation QteE exhibits on LasR when present, this antiactivator also targets RhlR when LasR is absent. In contrast, the antiactivation effect of QslA appears more specific to LasR activity, with only modest dampening of QS activity when LasR control is not present.

**Figure 1:**
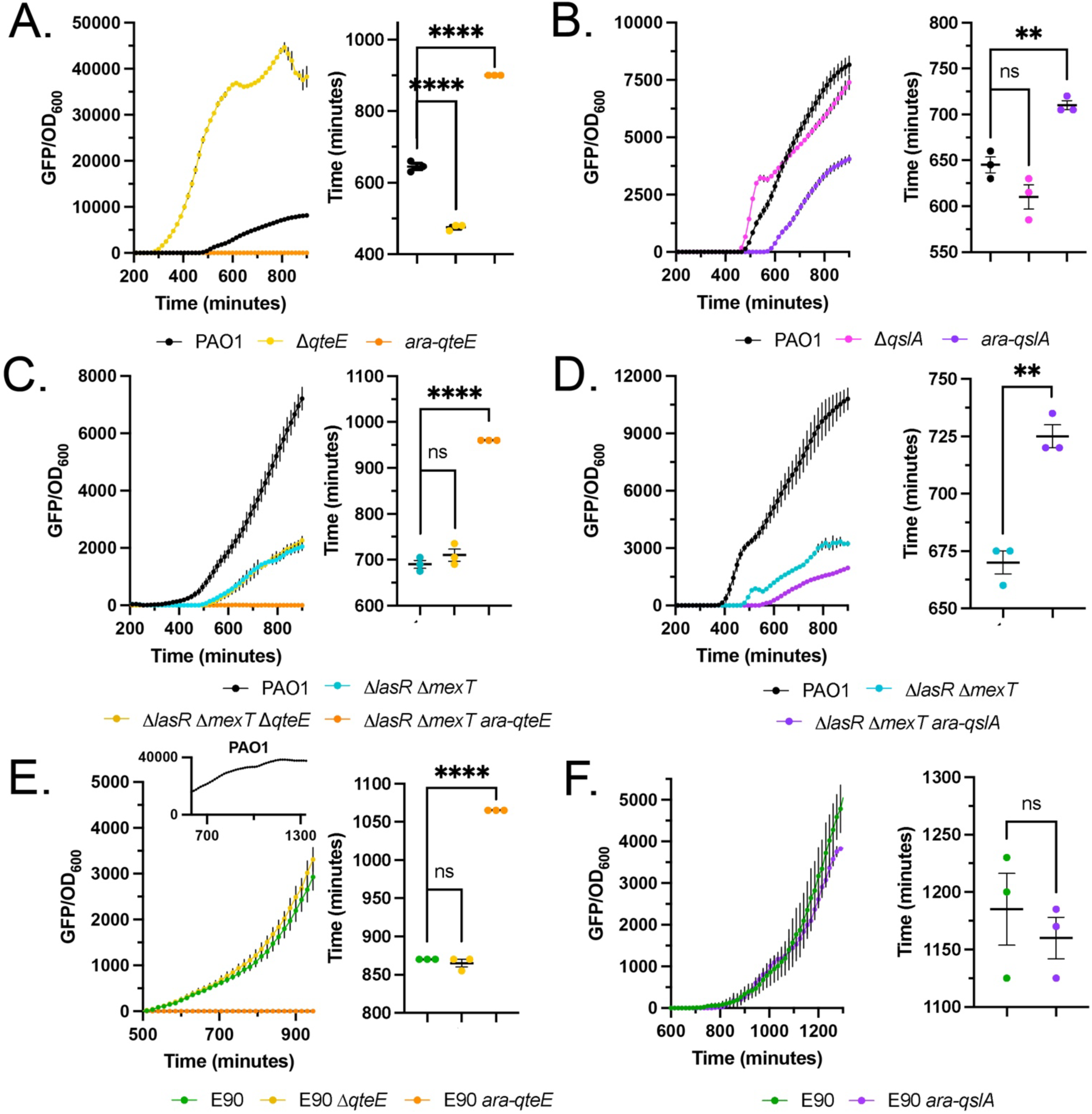
Effects of knocking out or over-expressing *qteE* (A, C, E) or *qslA* (B, C, F) on P*_rhlA_-gfp* expression over time in PAO1 (A, B), PAO Δ*lasR* Δ*mexT* (C, D), or E90 (E, F). Inset for (E) shows mean P*_rhlA_-gfp* expression in PAO1 in the same assay. *“ara*” indicates that the media was supplemented with 0.5% L-arabinose to induce over expression of the following gene. P*_rhlA_-gfp* activity was measured as GFP fluorescence and normalized to OD_600_ of experimental cultures. The y-axes for each plot are scaled to their maximal GFP levels. All data points show means of three biological replicates, with error bars representing s.e.m. Some error bars may not be visible. For each panel, fluorescence curves are on the left and time-to-half-maximal GFP measurements are on the right. Significant differences of means compared between indicated strains were determined using an unpaired two-tailed t-test (α = 0.05). ns, *p* > 0.05; **, *p* ≤ 0.01; **** *p* ≤ 0.0001.

Given our observations that over-expression of antiactivators delays or abolishes P*_rhlA_-gfp* induction in most cases, we questioned whether exogenous addition of C4-HSL could restore RhlR activity by increasing the availability of signal to bind and stabilize RhlR dimers. To test this idea, we repeated the over-expression experiments in PAO1 and PAO Δ*lasR* Δ*mexT* as detailed in Figure 1, but with supplementation of 10 μM C4-HSL. For this set of experiments, we also reduced the concentration of arabinose used (100-fold less for P*_araBAD_-qteE*, 10-fold less for P*_araBAD_-qslA*) to control antiactivator over-expression so we that we could obtain a non-zero measurements in the conditions without signal supplementation. We observed moderately restored P*_rhlA_-gfp* induction timing with C4-HSL supplementation in both PAO1 and PAO Δ*lasR* Δ*mexT* cells that over-express *qteE* (Figure 2A, 2C). A similar pattern is present in PAO1 and PAO Δ*lasR* Δ*mexT* cells that over-express *qslA*; supplementing these cultures with C4-HSL also advanced P*_rhlA_-gfp* induction timing, but to a greater degree than the experiments with *qteE* (Figure 2B, 2D). Regardless of the extent to which C4-HSL supplementation advanced timing or increased magnitude of P*_rhlA_-gfp* induction with respect to antiactivator deletions, over-expression of either antiactivator in either strain significantly reduced the time to half-maximal induction (Figure 2). Together, these results indicate QteE has RhlR-specific antiactivation activity independent of LasR status in a variety of strain backgrounds, while QslA activity appears to only affect RhlR activity in the laboratory strains and their mutants tested here.

**Figure 2:**
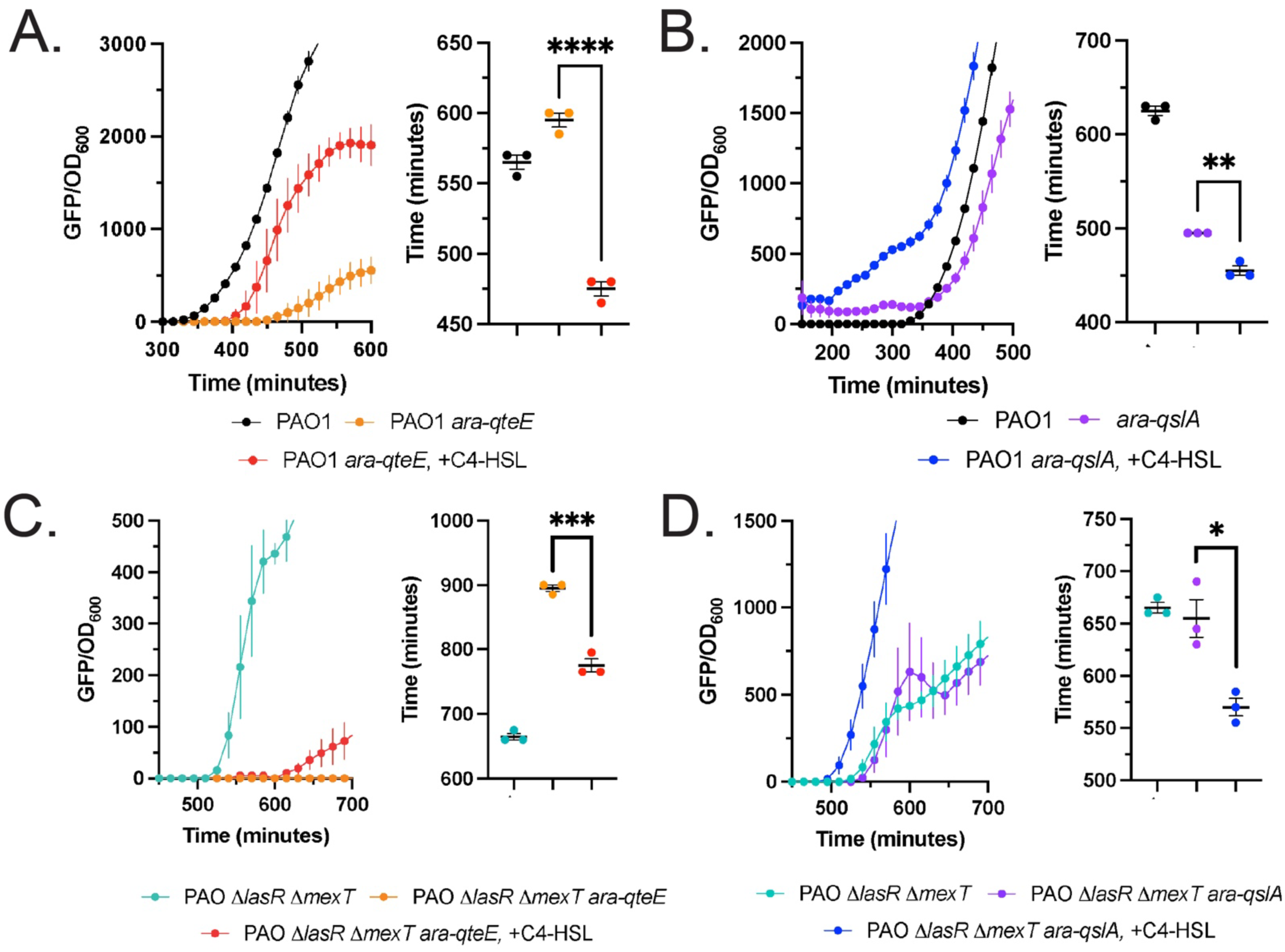
Effects of exogenous C4-HSL on PAO1 (A, C) or PAO Δ*lasR* Δ*mexT* (B, D) P*_rhlA_-gfp* expression with over-expressed *qteE* (A, B) or *qslA* (C, D). “*ara*” indicates that the media was supplemented with 0.005% L-arabinose for over-expressing *qteE* or 0.05% L-arabinose for over-expressing *qslA*. “+C4-HSL” indicates that the media was supplemented with 10 μM C4-HSL. P*_rhlA_-gfp* activity was measured as GFP fluorescence and normalized to OD_600_ of experimental cultures. Axes are scaled to focus on QS induction. All data points show means of three biological replicates, with error bars representing s.e.m. Some error bars may not be visible. For each panel, fluorescence curves are on the left and time-to-half-maximal GFP measurements are on the right. Significant differences of means compared between indicated strains were determined using an unpaired two-tailed t-test (α = 0.05). *, *p* ≤ 0.05; ***, *p* ≤ 0.001; ****, *p* ≤ 0.0001.

### Deleting *qslA* sensitizes LasR and RhlR to their signals

Given our observations that antiactivators alter RhlR QS induction timing, and that this effect can be C4-HSL signal concentration-dependent, we were curious if antiactivators could alter the threshold concentration for a cell population reach quorum (11, 15, 16). Previous literature has demonstrated that QteE slows the accumulation of both *P. aeruginosa* AHLs, and that deleting *qteE* triggers LasR activity (as measured by P*_lasI_* induction) at lower concentrations of 3OC12-HSL (11, 16). Therefore, we focused our experiments on QslA. To determine if QslA changes the concentration of 3OC12-HSL needed to trigger LasR induction, we introduced *lasI* and *qslA* deletion alleles into PAO1 and titrated 3OC12-HSL across a range of concentrations. We then measured LasR QS activity in batch cultures with a plasmid-borne GFP-transcriptional reporter constructed with the same backbone as P*_rhlA_-gfp,* but with the strictly LasR-specific promoter for *lasI* (P*_lasI_-gfp*). Similarly, to test the sensitivity of RhlR to C4-HSL in the context of QslA antiactivation, we introduced *rhlI* and *qslA* deletions into PAO Δ*lasR* Δ*mexT* and titrated C4-HSL across a range of concentrations, again using the plasmid-borne RhlR-specific P*_rhlA_-gfp* reporter.

In strain PAO Δ*lasI*, deletion of *qslA* results in significantly higher LasR induction at lower signal concentrations than in the parent strain. Supplementing growth media with just 10 nM 3OC12-HSL results in a 10-fold increase of P*_lasI_-gfp* in PAO Δ*lasI* Δ*qslA* compared to the *qslA-*proficient strain, nearly reaching maximal P*_lasI_-gfp* induction with the lowest supplement concentrations tested (Figure 3A). PAO Δ*lasI* Δ*qslA* also has a substantially lower saturating 3OC12 concentration than PAO Δ*lasI*; the *qslA*-deficient strain reaches peak P*_lasI_-gfp* expression at 100 nM 3OC12, while the parent strain reaches its maximum P*_lasI_-gfp* expression 200 nM, the highest concentration tested. Similar to the trend observed for QslA and LasR, deletion of *qslA* in PAO Δ*lasR* Δ*mexT* Δ*rhlI* results in significantly higher RhlR induction than in the parent strain at the tested C4-HSL concentrations, albeit to a lesser degree (Figure 3B). Notably, RhlR reporter activity in PAO Δ*lasR*Δ*mexT*Δ*rhlI* plateaus with additions of 5 μM C4-HSL or greater; in contrast, reporter activity in PAO Δ*lasR*Δ*mexT*Δ*rhlI*Δ*qslA* continually rises with higher concentrations of C4-HSL. We note that all non-zero signal concentration-matched differences between tested strains were significantly different. Taken together, results of these two experiments indicate that, similar to previous reports of QteE activity, QslA also alters QS induction thresholds in part by modulating sensitivity of LasR and RhlR to their cognate AHLs.

**Figure 3:**
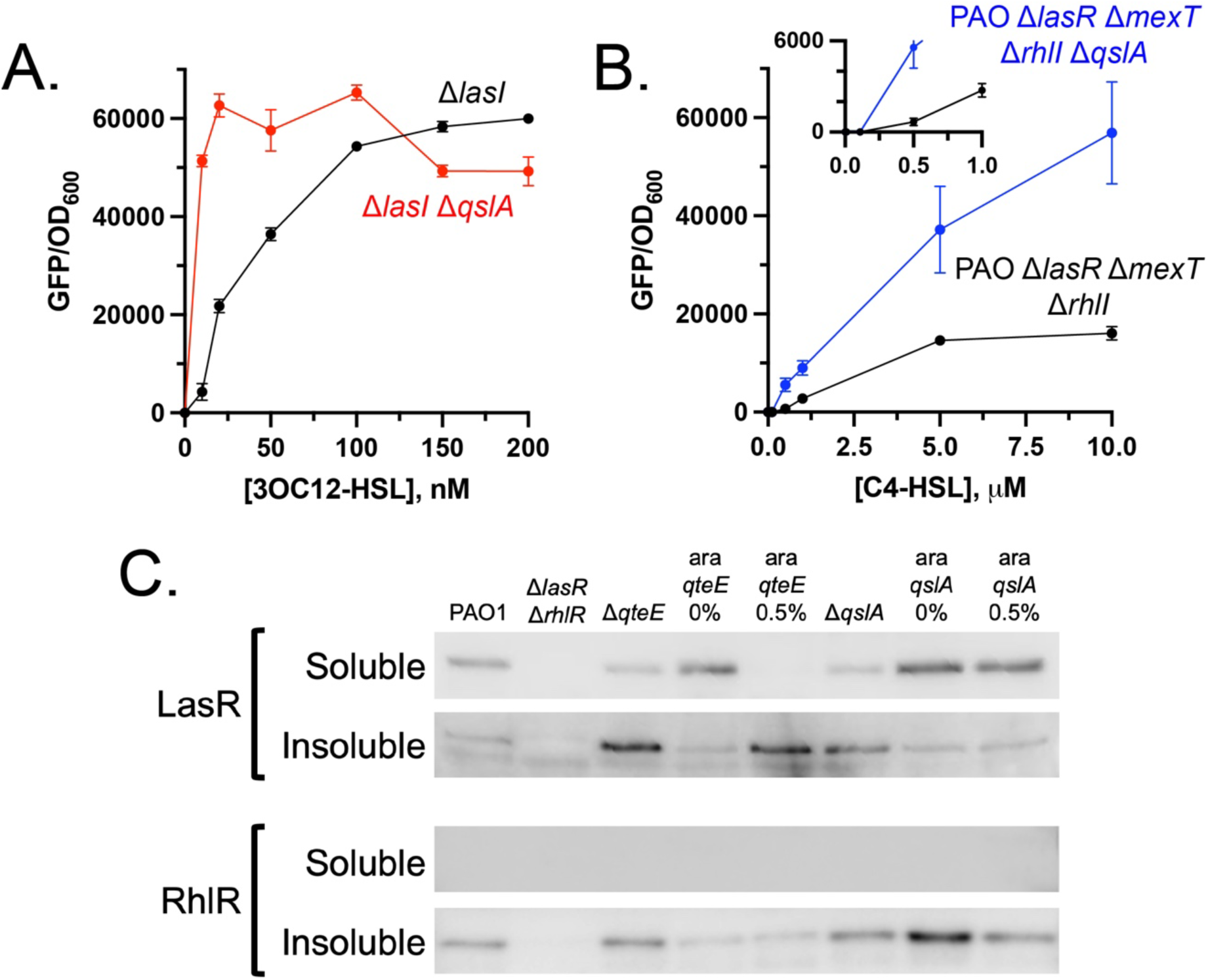
Effects of QteE or QslA on active and soluble LasR and RhlR. (A, B) Effects of QslA on sensitivity of LasR or RhlR to their cognate signals. (A) PAO1 strains were grown in varying concentrations of 3OC12-HSL. (B) PAO Δ*lasR* Δ*mexT* strains were grown in varying concentrations of C4-HSL. P_l_*_asI_-gfp* (A) or P*_rhlA_-gfp* (B) activity was measured as GFP fluorescence and normalized to OD_600_ of experimental cultures. Inset in (B) has reduced axes to emphasize reporter activity at low [C4-HSL]. Data points represent the mean of three biological replicates, and error bars represent s.e.m. Some error bars may not be visible. All non-zero signal concentration-matched differences in (A) and (B) were statistically significant, as assessed by two-tailed t-tests (α = 0.05). (C) LasR and RhlR Western blots. Samples were retrieved from cells harvested after 18 hours of growth (stationary phase). Percent values indicate the amount of L-arabinose supplemented for antiactivator over-expression. Roughly equal amounts of protein were added to each well. Bands from each row are from separate gels and therefore may not be comparable. Blots are representative of replicate experiments.

### QteE, but not QslA, decreases the amount of active and soluble LasR

Effects of antiactivators on AHL sensitivity could either be due to modification of R-protein-signal binding dynamics, or to direct modulation of active R-protein dimer levels through binding and sequestration. Given some previous evidence that QteE is known to destabilize LasR and RhlR dimers, and that QslA can bind LasR monomers to prevent LasR dimerization, we hypothesized the latter (11–13). To test this, we used Western blots to monitor the levels of LasR or RhlR in soluble and insoluble fractions of stationary phase culture cell lysates with deleted or over-expressed antiactivators (Figure 3C). Since active LasR and RhlR dimers are soluble in the cytoplasm while inactive LasR and RhlR monomers are not, measuring soluble and insoluble fractions separately provides a proxy for the levels of active and inactive R-protein dimers and monomers, respectively.

Over-expressing *qteE* removes any LasR from the soluble fraction and dramatically increases levels of LasR present in the insoluble fraction (Figure 3C). In contrast, over-expressing *qteE* does not appear to markedly change RhlR levels in the insoluble fraction. No soluble RhlR can be observed in these blots, as is consistent with previous literature noting a lack of RhlR solubility in the absence of C4-HSL, but some insight may be gleaned from data on the insoluble fraction (32, 33). These results indicate QteE largely functions by destabilizing soluble and active LasR dimers, as over-expressing *qteE* appears to force most LasR protein into the insoluble fraction. Interestingly, over-expressing *qteE* does not change the levels of RhlR in the insoluble fraction but deleting *qteE* appears to increase this fraction slightly relative to WT. On the other hand, over-expression of *qslA* has no marked effect on LasR levels or solubility. Over-expression of *qslA* does appear to change RhlR levels in the insoluble fraction, but it is unclear if this effect is independent of QslA effects on LasR. Overall, these results indicate that while QteE appears to decrease the amounts of active and soluble LasR and RhlR, QslA-mediated antiactivation may work via a different inhibitory mechanism.

### QteE, but not QslA, is regulated by LasR and RhlR

As previous data has shown LasR and RhlR exhibit strong QS-regulated autoinduction, we considered that QteE and QslA, as factors important in QS activation, may also be transcriptionally regulated by QS. Previous microarray studies in PAO1 have suggested that *qteE* transcription may be repressed by QS under some conditions, but this result has not been substantiated in followup studies (7, 15). To clarify the individual roles of LasR and RhlR on antiactivator regulation, we utilized qRT-PCR to measure mRNA transcript levels from either *qteE* or *qslA* in stationary phase cultures of PAO1, PAO Δ*lasR*, and PAO Δ*rhlR* (OD_600_ = 2.0). We observed higher *qteE* ΔC_t_ values in PAO Δ*lasR* and PAO Δ*rhlR* when compared to PAO1 (Figure 4A), indicating both R-proteins repress QteE transcription. In contrast, the absence of LasR or RhlR does not change *qslA* transcription significantly (Figure 4B). These results confirm a role for QS regulation of *qteE*, but not *qslA*.

**Figure 4:**
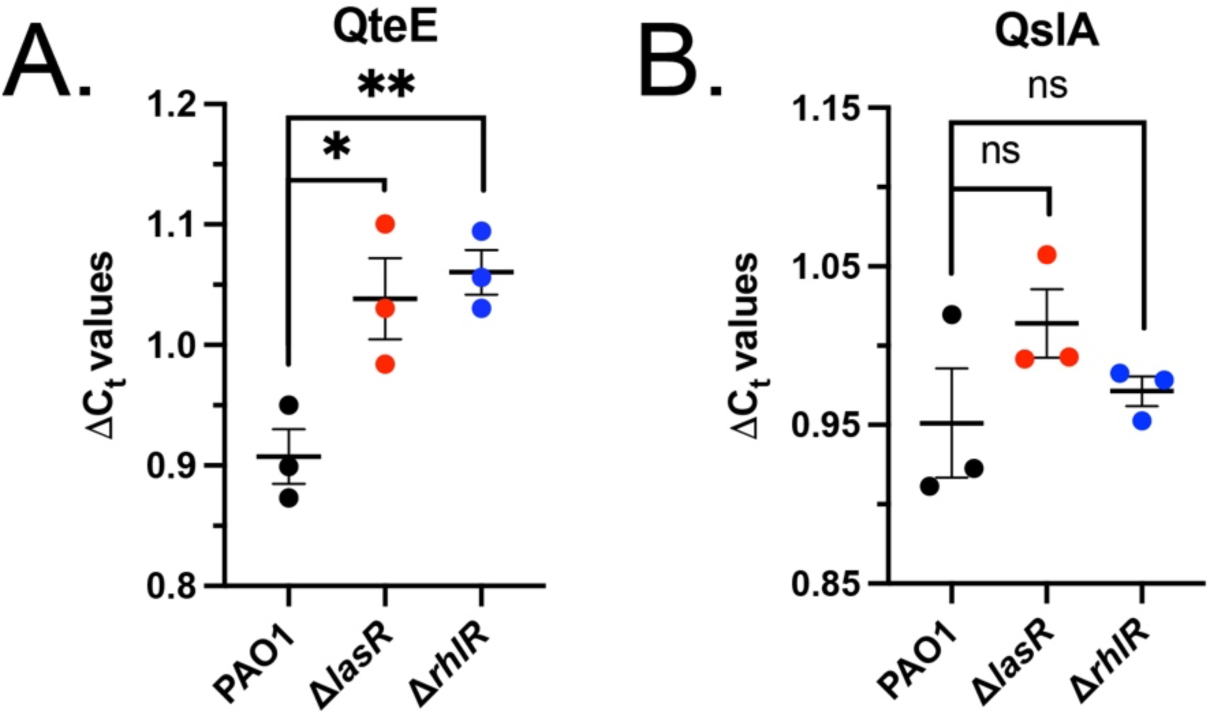
QS regulation of QteE or QslA. Expression of *qteE* (A) and *qslA* (B) in PAO1, PAO Δ*lasR*, and PAO Δ*rhlR* were measured with qRT-PCR. Differences in threshold cycle (ΔC_t_) values are normalized to the housekeeping gene *rplU*. Data points represent biological replicates, with means and error bars indicating s.e.m. Significant differences of means compared between indicated strains were determined using an unpaired two-tailed t-test (α = 0.05). ns, *p* > 0.05; *, *p* ≤ 0.05; ***, *p* ≤ 0.001.

### QslA effects on QS phenotypes may depend on QteE

Our results and previous literature indicate that QteE may exert its regulatory activity at lower cell densities and earlier in the QS cascade than QslA in PAO1, presenting the possibility that QS phenotypes delayed by the latter depend on the presence of QteE (11–13). Analysis of single and multiple antiactivator deletions in PAO1 suggest their effects on QS induction can be additive, but the dependence of QslA activity on *qteE* status has not been explored (15). We hypothesized that in the absence of QteE, the dampening effect of QslA on QS regulation and phenotypes would be reduced. We tested this idea by measuring two quorum-controlled virulence phenotypes in stationary phase cultures of a panel of PAO1-derivative mutants. Because the regulatory dependence in question would require a strain background capable of both LasR and RhlR QS, we used the PAO1 lab strain and analyzed phenotypes that are predominantly regulated by LasR (elastase) or RhlR (pyocyanin) in this lineage. We again used antiactivator deletion mutants and our separate chromosomal P*_araBAD_-qteE* and P*_araBAD_-qslA* over-expression constructs with arabinose supplementation to control antiactivation. As previous studies have demonstrated, deletions of *qteE* or *qslA* alone results in increases in both elastase activity and pyocyanin production (Figure 5). Over-expressing *qteE* in PAO1 and PAO Δ*qslA* reveals complete abrogation of both elastase activity and pyocyanin production, equivalent to a QS-silent Δ*lasR* Δ*rhlR* mutant. Consistent with our hypothesis, over-expressing *qslA* in PAO1 reduces elastase activity (although not significant (p = 0.2185)) but doing so in PAO Δ*qteE* has no notable effect on elastase activity (Figure 5A). *qslA* over-expression does slightly reduce pyocyanin production in the context of a Δ*qteE* deletion mutant, but not significantly in PAO1 (Figure 5B).

**Figure 5:**
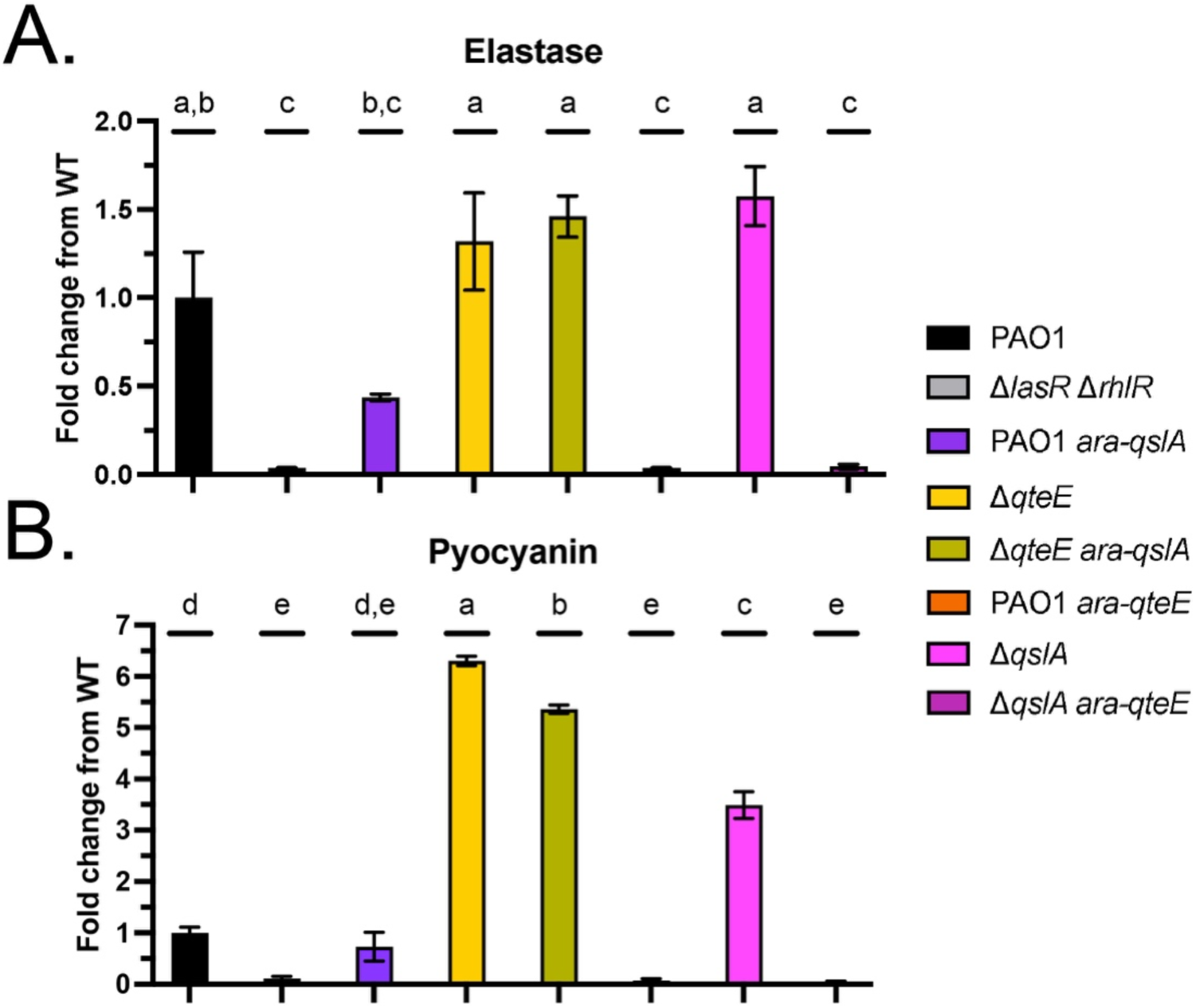
Antiactivator interdependency in QS-controlled phenotypes. (A) elastase activity and (B) pyocyanin production. Each assay was performed after 18 hours of growth (stationary phase cultures). “*ara-qslA*” and “*ara-qteE*” indicate the presence of an arabinose-inducible construct in the chromosome, and the culture was supplemented with 0.5% L-arabinose to trigger over-expression of the corresponding antiactivator. Values represent the mean of three biological replicates, and error bars represent s.e.m. Letters indicate statistically significant differences of means, as determined using an ordinary one-way ANOVA.

## Discussion

*P. aeruginosa* uses QS to control gene expression underlying several important phenotypes and population-level behaviors, aiding this bacterium in nutrient acquisition, survival during stress, and virulence during infection (34). The effectiveness of *P. aeruginosa* QS in yielding these phenotypes is in part due to factors that tightly control circuit activation and target gene transcription, including several mechanisms that serve to dampen QS (3, 11, 12, 35). Here, we focused on so-called “antiactivators” of QS: protein factors that interact directly with R-proteins to attenuate activation in populations below quorum. Our results support a model where antiactivation dynamics in *P. aeruginosa* are nuanced in both the specificity of antiactivator-LuxR-protein interactions, as well as the magnitude of their effects on QS activation.

Separating the effects of antiactivation on RhlR from those on LasR has proved challenging due to the hierarchy of QS in laboratory strains. Furthermore, previous antiactivator studies that assess the kinetics of R-protein induction have measured *lasB* expression, as there is strong QS activation of this gene. The demonstrated promiscuity of the *lasB* promoter to allow both LasR and RhlR binding has made it difficult to dissect differing effects of antiactivation on the two regulons (15, 36). This has a left a notable knowledge gap regarding the effects of antiactivation on RhlR QS. To address this gap, we analyzed the laboratory strain mutant PAO Δ*lasR* Δ*mexT* and clinical isolate E90, both demonstrating LasR-independent RhlR QS, alongside PAO1 to parse the individual effects of antiactivators on RhlR activity. Our results show a strong QteE-mediated antiactivation effect on RhlR in a variety of circumstances. Over-expressing *qteE* not only delays RhlR induction but completely abrogates this activity in all strains tested regardless of the LasR status of the strain. The strength of QteE antiactivation on RhlR was strong enough that even a 100-fold reduction in the typical concentration of arabinose used to induce our synthetic construct still produced significant dampening of target gene activation. It is then notable that deleting *qteE* in the two strains without LasR, PAO Δ*lasR* Δ*mexT* and E90, has no effect on P*_rhlA_-gfp* induction. We recognize that *qteE* may not be well-expressed in the strains with RhlR*-*dominated QS that we evaluated or may not be expressed in the conditions tested here.

In contrast to QteE, over-expressing *qslA* only influenced RhlR activity in PAO1 and PAO Δ*lasR* Δ*mexT*, having no marked effect on RhlR activity in E90. It is intriguing that QslA modulates RhlR induction timing differently in two strain backgrounds exhibiting Las-independent QS. Expression of some RhlR-regulated genes is rescued in PAO Δ*lasR* Δ*mexT* via another QS pathway, the Pseudomonas quinolone signal (PQS) system (28, 37). QslA could be interacting with the PQS receptor-regulator PqsR in PAO Δ*lasR* Δ*mexT*; such an interaction has been demonstrated in the rhizosphere isolate PA1201 (38). Confirmation of this interaction in future studies would add another layer of complexity to the QS dynamics in some of the most widely-used laboratory strains of *P. aeruginosa.* Further investigation of the genomics and QS signaling diversity of additional *P. aeruginosa* clinical isolates may also further define the role of QslA in strains lacking LasR.

We observe that QslA expression, as seen in previous studies of QteE, decreases the sensitivity of LasR and RhlR to their cognate signals (11, 16). A potential explanation is that without these antiactivators to sequester inactive LasR and RhlR monomers, any R-protein that is produced can bind its cognate signal and become active without further inhibition, allowing a cell to short-circuit QS and “sense itself” regardless of cell and signal density. Careful experimentation by others at extremely low cell densities has revealed this to be true for the *las* system in subpopulations of cells, where some LasR gene targets exhibit expression that appears constitutive due to “self-sensing” in the absence of antiactivation (16). Indeed, we find that addition of the LasR-specific 3OC12-HSL signal to a Δ*lasI* Δ*qslA* mutant of PAO1 dramatically advances the induction of a LasR-specific promoter, even at levels 20-fold less than maximal WT signal production. A similar effect is observed with additions of the RhlR-specific C4-HSL signal to a Δ*lasR* Δ*mexT* Δ*rhlI* Δ*qslA* mutant, although the effect is subdued in comparison the *las* system. This may reflect a nuance with how RhlR and its cognate signal interact, as compared to most LuxR homologs: in strain PAO1, RhlR exhibits signal sensitivity and binding kinetics that are vastly different from most well-studied QS receptors, with the C4-HSL concentration required for half-maximal activity 50-times higher than any other LuxR protein analyzed (39). Our results expand the role of antiactivators in preventing self-sensing through LasR; QteE and QslA may also prevent RhlR self-sensing in strains that lack LasR.

Our Western analysis shows that over-expression of *qteE* causes a complete loss of LasR in the soluble fraction and a substantive increase of LasR in the insoluble fraction. This corroborates previous literature that shows inducing *qteE* expression destabilizes LasR, and the increased insoluble LasR is likely targeted for degradation (11). Notably, over-expressing *qteE* has no effect on the level of RhlR in the insoluble fraction, but deleting it raises it. A potential explanation is that leakage from the P*_araBAD_* promoter used to trigger over-expression allows sufficient QteE production to attenuate RhlR at a saturating level. While previous work already showed that QteE destabilizes LasR proteins in whole cell lysates, here we provide additional evidence that QteE-mediated destablization may function by forcing inactive LasR monomers into the insoluble fraction (11). We also recognize that further investigation is needed to find an approach for analyzing RhlR stability in conditions where it retains more solubility, as the overall lack of RhlR solubility in lysates has been observed in other Western analyses. Availability of the C4-HSL signal is required for RhlR binding of target promoters, and addressing the dependence of RhlR solubility on C4-HSL may provide a pathway forward (5). On the other hand, over-expressing *qslA* does not markedly change soluble or insoluble levels of LasR; given some evidence that QslA binds inactive LasR monomers to prevent dimerization without necessarily targeting them for degradation, this is not surprising (12, 13). While over-expressing *qslA* does change RhlR levels in the insoluble fraction, this may be due to a downstream effect of QslA attenuating LasR, and RhlR solubility issues may also complicate this interpretation.

We sought to address the potential separate effects, if any, of QS regulation via LasR or RhlR on antiactivator expression. Our expression analysis in batch cultures demonstrate that LasR appears to repress transcription of *qteE*, but not *qslA*. The finding that *qteE* is repressed by LasR is in agreement with some previously published RNA transcriptome comparisons of PAO1 and a QS-null mutant (7, 15, 40). While Schuster *et al*. reported a 29-fold increase in *qteE* expression in an R-protein-deficient strain compared to WT PAO1 when both tested strains were grown in a rich media, two additional studies reported no change in *qteE* expression when the same strains were tested in more defined media (7, 15, 40). Given that each study was performed in different strain backgrounds, and with different media and growth phases, the contrast is somewhat unsurprising. The possibility that QteE is indirectly regulated by LasR and RhlR could also explain these differences and cannot be ruled out based on our data. However, a repressive effect of QS on the transcription and availability of QteE could provide *P. aeruginosa* further means for reinforcing an activated QS state by effectively silencing this mode of antiactivation. Our observation that RhlR is also regulated this way provides further strength for this interpretation. In this view, QteE acts to reinforce the bistability in *P. aeruginosa* QS response: while QteE is available to bind LasR or RhlR and help keep QS activation low in concordance with cell density, as the signal threshold is reached and QS becomes activated, QteE is in turn repressed to support the QS-active state. QslA does not appear to be QS-controlled, on the other hand. It is possible QslA may instead provide a more stable dampening effect on QS that is overcome upon activation, or this could be due to a general lack in native expression in the conditions of our experiments. The role of antiactivation in influencing heterogeneity in the QS response, particularly at the single-cell level, is a related area poised for active investigation. Heterogeneity in response between individual cells in a population is increasingly recognized as general feature of QS systems, and examples have been presented for both early and transient heterogeneity in the early phases of QS activation, as well as persistent heterogeneity through high cell densities and activation states (41–44). A recent study of the impacts of another negative regulator of QS, the transcriptional repressor of *lasI/lasR,* RsaL, revealed this repressor provides a degree of stochasticity in single-cell responses within the *las* system as a quorum threshold is approached (45). The same study reported greater homogeneity in responses in the *rhl* QS system, which lacks a transcriptional repressive mechanism analogous to RsaL. Further investigation of divergent QS systems lacking the RsaL-*las* dynamic, coupled with dissection of the differential regulation and effects of individual antiactivators, should offer utility to future studies of the origins and impacts of QS heterogeneity.

We also find that QslA-mediated effects on quorum-controlled phenotypes show some dependence on QteE. Existing structural evidence combined with the lack of similarity between QteE and QslA sequences suggest antiactivators interact with R-proteins separately, yet the layered nature of QS regulation in *P. aeruginosa* leaves several opportunities for dependent effects. QteE acts to potently silence LasR early in the growth cycle of PAO1, so we expected to observe muted effects of QslA on downstream phenotypes owing to the QS hierarchy present in this strain. However, this effect was only true for elastase production: deletion of *qteE* does not remove the impact of excess QslA on pyocyanin production. We acknowledge that the magnitude of this effect could be due to levels of QslA not physiologically relevant to native QS systems, but the observed impact could be instructive. There is also reason to suspect nutrient or growth status may dictate how antiactivators affect the differential regulation of specific QS-controlled secretions. The LasR and RhlR regulators are embedded in a network of other regulatory components that push and pull on the activation threshold to tailor the QS response beyond just cell and signal density (46). The availability of specific macronutrients, for example, can alter the levels of corresponding QS-controlled secretions that contain the nutrient, likely via the stringent response (47). Pyocyanin production involves a complex phenazine biosynthesis pathway with additional inputs to regulation, particularly through regulators of the PQS system, which may also interact with QslA. Further investigation of the links between the *rhl* QS and PQS systems may provide further insight in this regard, as will definition of the response to physiological status by antiactivation mechanisms.

Our results contribute additional detail to a general understanding of the complex circuitry of *P. aeruginosa* QS, underscoring a distinct role for QteE in the repression of RhlR QS activity. We demonstrate the differing mechanisms and regulation of QteE and QslA can have distinct consequences for LasR and RhlR function as cells transition from low to high cell densities, and that regulation of these mechanisms is also variable among divergent QS circuits. Our study illustrates that antiactivation is not a single repressive mechanism accomplished by several factors, rather individual antiactivators may have plenary effects on QS (QteE) or more targeted effects on specific R-proteins (QslA), providing finer tuning of QS activation for varying physiological states.

## Materials and Methods

### Culture conditions

All strains and experimental cultures were grown at 37°C either on LB agar or in LB broth buffered with 50 mM 3-(*N*-morpholino)-propanesulfonic acid (MOPS), pH = 7.0, with shaking at 250 rpm. When necessary, growth media was supplemented with 100 μg/mL gentamicin or 200 μg/mL carbenicillin to select for marked *P. aeruginosa* strains, and 10 μg/mL gentamicin or 200 μg/mL carbenicillin to select for *E. coli* strains carrying plasmids. Reporter plasmids were maintained in *P. aeruginosa* cultures by growing strains in media supplemented with 100 μg/mL gentamicin. All strains assessed in Figures 1 and 2 grew normally (Figures S3 and S4).

### Strain and plasmid construction

A full list of the strains used in this study is detailed in Table 1. *P. aeruginosa* PAO1 (UW) or the EPIC isolate E90 were used as the parent strain for all strain construction (48). In all experiments, PAO1 was used as the wild-type control strain. PAO1, Δ*lasR*, Δ*rhlR,*Δ*lasR*Δ*rhlR*, and E90 were obtained from the Dandekar Lab at the University of Washington. PAO Δ*qteE* was obtained from R. Siehnel (11). The PAO Δ*qslA*, Δ*lasI*, and Δ*rhlI* deletion alleles were constructed using a pEXG2-based suicide delivery vector carried in *Escherichia* coli S17 following a two-step allelic exchange protocol (49). A pEXG2-Δ*mexT* plasmid was obtained from M. Kostylev (28). The deletion allele for all mutants consists of the first two residues and the last two residues of the protein of interest. Strains with over-expressed *qteE* or *qslA* were constructed by inserting an extra copy of the corresponding gene under P*_araBAD_* control (arabinose-inducible) into the *P. aeruginosa* chromosome at a neutral site using a mini-Tn7 mediated integration (50). Briefly, we cloned the mini-Tn7 machinery, the P*_araBAD_* sequence, and the CDS of either *qteE* or *qslA* into pUC19T using a modified Gibson Assembly (51, 52). The Gm^R^ marker on this plasmid was removed from successful transposon mutants using a Flp-mediated excision (50). In experimental cultures, over-expression was triggered by supplementing growth media with 0.5% arabinose unless stated otherwise. pPROBE-GT transcriptional reporters for *lasI* and *rhlA* were obtained from the Dandekar Lab.

**Table 1:** List of strains used in this study.

| Strain | Relevant information | Origin |
| --- | --- | --- |
| <u>*P. aeruginosa* strains</u> |  |  |
| PAO1 (UW) | Wild-type, UW library strain | Stover *et al.*, 2000 |
| PAO $\Delta qteE$ | PAO1 derivative, markerless *qteE* deletion mutant | Siehnal *et al.*, 2010 |
| PAO1 $P_{araBAD}$-*qteE* | PAO1 derivative, extra copy of *qteE* under $P_{araBAD}$ control at *attTn7* site | This study |
| PAO $\Delta qslA$ | PAO1 derivative, markerless *qslA* deletion mutant | This study |
| PAO1 $P_{araBAD}$-*qslA* | PAO1 derivative, extra copy of *qslA* under $P_{araBAD}$ control at *attTn7* site | This study |
| PAO $\Delta lasR \Delta mexT$ | PAO1 derivative, markerless *mexT* deletion in a *lasR* mutant | Kostylev *et al.*, 2019 |
| PAO $\Delta lasR \Delta mexT \Delta qteE$ | PAO $\Delta lasR \Delta mexT$ derivative, markerless *qteE* deletion mutant | This study |
| PAO $\Delta lasR \Delta mexT P_{araBAD}$-*qteE* | PAO $\Delta lasR \Delta mexT$ derivative, extra copy of *qteE* under $P_{araBAD}$ control at *attTn7* site | This study |
| PAO $\Delta lasR \Delta mexT P_{araBAD}$-*qslA* | PAO $\Delta lasR \Delta mexT$ derivative, extra copy of *qslA* under $P_{araBAD}$ control at *attTn7* site | This study |
| E90 | Chronic infection isolate from pediatric CF patient | Feltner *et al.*, 2016 |
| E90 $\Delta qteE$ | E90 derivative, markerless *qteE* deletion | This study |
| E90 $P_{araBAD}$-*qteE* | E90 derivative, extra copy of *qteE* under $P_{araBAD}$ control at *attTn7* site | This study |
| E90 $P_{araBAD}$-*qslA* | E90 derivative, extra copy of *qslA* under $P_{araBAD}$ control at *attTn7* site | This study |
| PAO $\Delta lasI$ | PAO1 derivative, markerless *lasI* deletion mutant | This study |
| PAO $\Delta lasI \Delta qslA$ | PAO1 derivative, markerless *lasI* and *qslA* deletion mutant | This study |
| PAO $\Delta lasR \Delta mexT \Delta rhII$ | PAO $\Delta lasR \Delta mexT$ derivative, markerless *rhII* deletion mutant | This study |
| PAO $\Delta lasR \Delta mexT \Delta rhII \Delta qslA$ | PAO $\Delta lasR \Delta mexT$ derivative, markerless *qslA* and *rhII* deletion mutant | This study |
| PAO $\Delta lasR \Delta rhIR$ | PAO1 derivative, markerless *lasR* and *rhIR* deletion double mutant | Siehnal *et al.*, 2010 |
| PAO $\Delta qteE P_{araBAD}$-*qslA* | PAO $\Delta qteE$ derivative, extra copy of *qslA* under $P_{araBAD}$ control at *attTn7* site | This study |
| PAO $\Delta qslA P_{araBAD}$-*qteE* | PAO $\Delta qslA$ derivative, extra copy of *qteE* under $P_{araBAD}$ control at *attTn7* site | This study |
| PAO $\Delta lasR$ | PAO1 derivative, markerless *lasR* deletion mutant | Wang *et al.*, 2015 |
| PAO $\Delta rhIR$ | PAO1 derivative, markerless *rhIR* deletion mutant | Wang *et al.*, 2015 |
| <u>*E. coli* strains</u> |  |  |
| *E. coli* DH5α | F- $\Phi 80 lacZ \Delta M15 \Delta(lacZYA-argF)$ U169 *recA1 endA1 hsdR17* (rk-, mk+) *phoA supE44* λ-*thi-1 gyrA96 relA1* | NEB |
| S17-1 | *recA pro hsdR* RP4-2-Tc::Mu-Km::Tn7 | Simon *et al.*, 1983 |

### GFP Transcriptional Reporter Assays

Transcriptional reporter assays were performed using a GFP promoter fusion of *lasI* or *rhlA* carried in pPROBE-GT. 18-hour endpoint assays were performed by inoculating experimental strains into 3 mL LB+MOPS, incubating with shaking for 18 hours, then measuring cell density (OD_600_) and GFP fluorescence (λ_excitation_ = 480 nm, λ_emission_ = 535 nm, gain = 80). Data is reported as a ratio of GFP fluorescence to cell density. GFP time course assays were performed as previously described (53). Briefly, experimental strains were inoculated at OD_600_ = 0.01 into 200 μL LB+MOPS in a black-walled 96-well plate which was incubated at 37°C with shaking. Cell density and fluorescence were measured at 15-minute intervals for 15 hours in a Biotek Synergy H1 microplate reader. GFP reporter activity for each timepoint was normalized to OD_600_ of the corresponding strain, and experimental GFP/OD_600_ values were corrected by subtraction from an empty promoter-less control pPROBE-GT plasmid. When performing experiments to assess *qslA* over-expression in E90, the 96-well plate was covered with a Breath Easy sealing membrane and incubated at 37°C with shaking for 7 hours in a standard shaking incubator before being measured in a microplate reader as described above. Statistical analysis of P*_rhlA_-gfp* induction timing was performed in GraphPad Prism by measuring time to half-maximal GFP for each replicate and performing an unpaired two-tailed t-test to compare strains of interest (α = 0.05). Growth curves of all strains analyzed were inspected to confirm no differences in growth dynamics were present (Supplemental Figures S3, S4).

### Western Blots

Cell fractionation as well as the assessment of insoluble and soluble LasR and RhlR levels were performed as per a previously described protocol (3). Briefly, experimental strains were inoculated into 3 mL LB+MOPS and were incubated at 37°C with shaking for 18 hours. Cells were pelleted and resuspended in LasR Purification Buffer (25 mM Tris-HCL [pH 7.8], 150 mM NaCl, 1 mM dithiothreitol [DTT], 1 mM EDTA, 10% glycerol, 0.05% Tween 20) with added 2 μM 3OC12-HSL and 10 μM C4-HSL. Samples were set to the same OD_600_ upon resuspension of the pellets to normalize the total protein content in each sample. Soluble and insoluble fractions were separated by sonicating samples and centrifuging at 55,000 g for 30 minutes at 4°C. The soluble fraction (supernatant) was removed, and the insoluble fraction (pellet) was resuspended in the same volume of LasR Purification Buffer with AHLs. 10 μL of the soluble and insoluble fractions were separated on a 4-12% Bis-Tris gel (Invitrogen). Separated proteins were transferred to a PVDF membrane and probed with polyclonal anti-LasR and anti-RhlR antibodies (3), then visualized using chemiluminescence detection reagents (Thermo Fisher Scientific).

### RNA extraction and qRT-PCR

Experimental strains were inoculated into 25 mL LB+MOPS at OD_600_ = 0.005 and incubated at 37°C with shaking in 125-mL baffled flasks. 1 mL of cells were harvested once the cultures reached OD_600_ = 2.0 and immediately stabilized with RNAprotect Bacteria Reagent (Qiagen). The cells were then pelleted and frozen at -80°C until RNA extraction. Pellets were resuspended in QIAzol reagent (Qiagen) and mechanically lysed by bead-beating. RNA extraction was performed using column-based purification via a RNeasy Mini Kit (Qiagen). RNA samples were then treated with Turbo DNase (Invitrogen) and purified with a RNeasy MinElute Cleanup Kit (Qiagen). cDNA was synthesized using an iScript cDNA Synthesis Kit (Bio-Rad). Gene expression was measured using iQ SYBR Green SuperMix (Bio-Rad) on a CF96X real-time PCR thermocycler for 40 cycles. The housekeeping gene *rplU* was used as a reference gene. A list of primers used for qRT-PCR is detailed in Table 2. Statistical analysis was performed using a two-tailed unpaired t-test in Graphpad Prism to compare means of expression levels between strains.

**Table 2:**
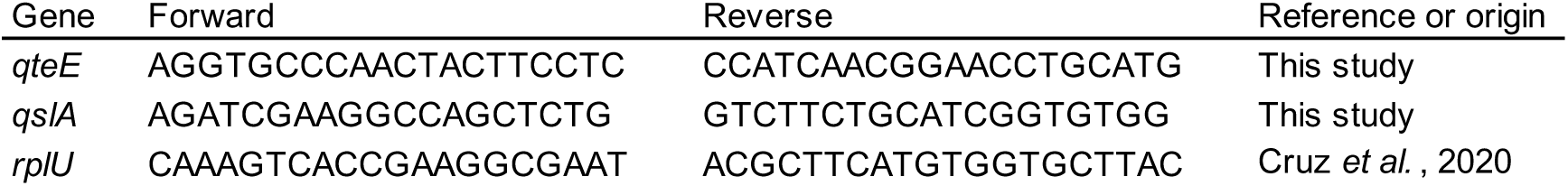
List of primers used for qRT-PCR in this study.

### Elastase activity assays

Elastase activity was measured using a variation on an elastin congo red (ECR) assay that allows for high throughput, as previously described (15, 54). Briefly, experimental strains were inoculated into 800 μL LB+MOPS in 96-well deep-well blocks at OD_600_ = 0.01 that were covered with Breath Easy sealing membranes and incubated at 37°C with shaking. After 18 hours, cells were pelleted at 4000 rpm for 10 minutes. 250 μL supernatant was filtered using 96-well filter plates, and 40 μL filtered supernatant was added to 360 μL ECR buffer (100 mM Tris, 1 mM CaCl_2_, pH = 7.5) with 20 mg/mL ECR. Samples were incubated with ECR in 96-well deep-well blocks and incubated at 37°C with shaking for 3 hours. Insoluble ECR was pelleted at 4000 rpm for 10 minutes, and 200 μL supernatant was transferred to 96-well plates and measured at ABS_495_ with a microplate reader. Final absorbance values were normalized to OD_600_ of the experimental cultures, and data is reported as fold change from WT samples. Statistical analysis was done in GraphPad Prism by performing an ordinary one-way ANOVA between all strains.

### Pyocyanin production assays

Pyocyanin production was measured using a previously described procedure (55). Briefly, experimental strains were inoculated into 5 mL LB+MOPS at OD_600_ = 0.01 in disposable 14 mL culture tubes (Falcon) and incubated at 37°C with shaking for 18 hours. Pyocyanin was extracted from the cultures using 3 mL chloroform, then extracted again from the organic phase with 660 μL 0.2 M HCl. 200 μL of the aqueous phase containing acidified pyocyanin was measured at ABS_520_ with a microplate reader. Final absorbance values were normalized to OD_600_ of the experimental cultures, and data is reported as fold change from WT samples. Statistical analysis was done in Graphpad Prism by performing an ordinary one-way ANOVA between all strains.

## Data availability

All data used in this study is contained herein.

## Acknowledgments

V.S. was funded in part by the CFF Student Traineeship Award SRIDHA20H0 and the Levinson Emerging Scholar Award at the University of Washington - Seattle. K.L.A was funded by CFF Fellowship ASFAHL19F0. A.A.D. was supported by NIH grants R01GM125714 and R35GM152107.

---

**Figure S1:**
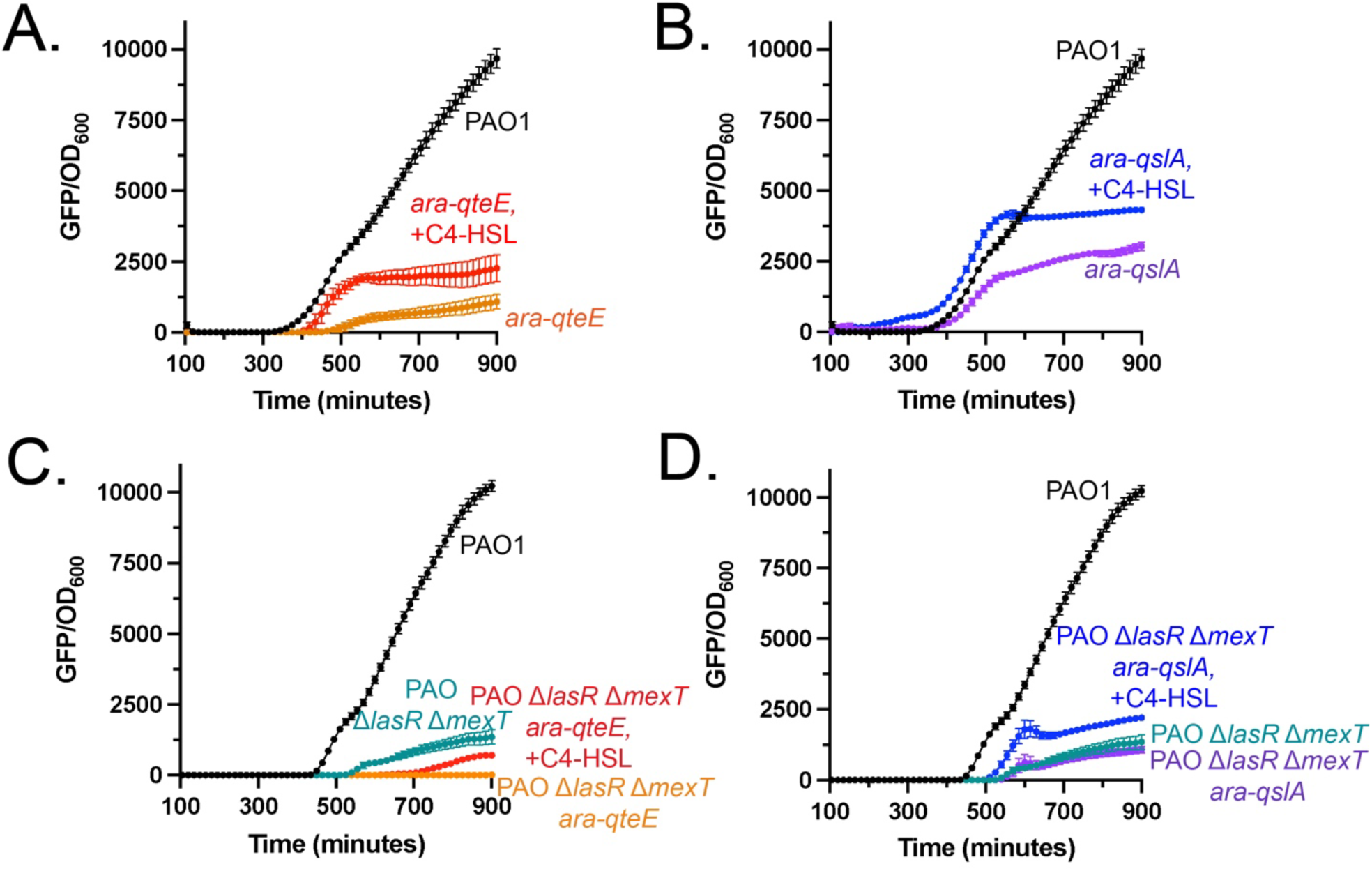
Uncropped plots of those shown in Figure 2. All data points show means of three biological replicates, and error bars represent s.e.m. Some error bars may not be visible.

**Figure S2:**
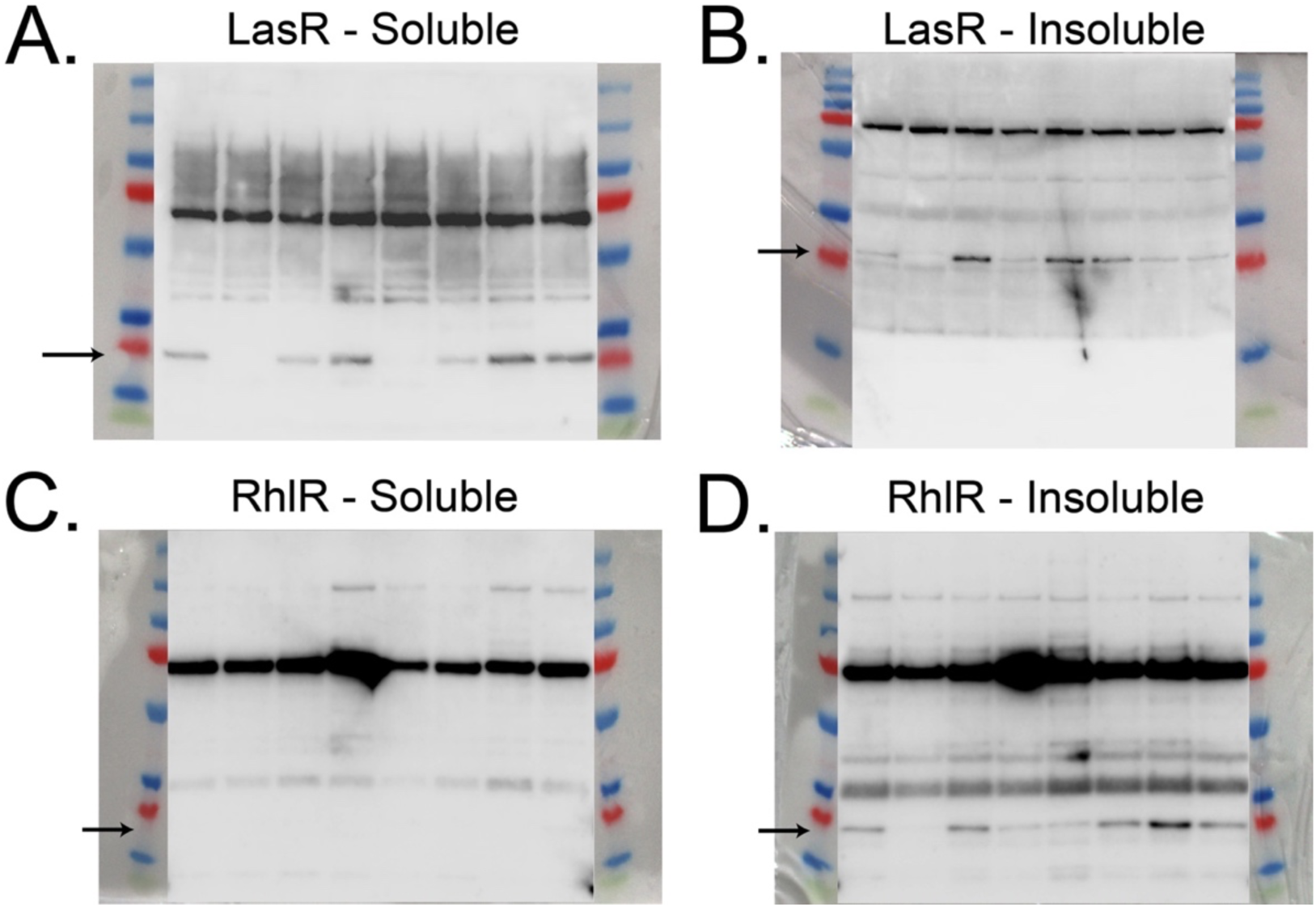
Uncropped blots of those shown in Figure 3. Expected LasR (A, B) or RhlR (C, D) bands are indicated by arrows. Blots are indicative of replicated experiments.

**Figure S3:**
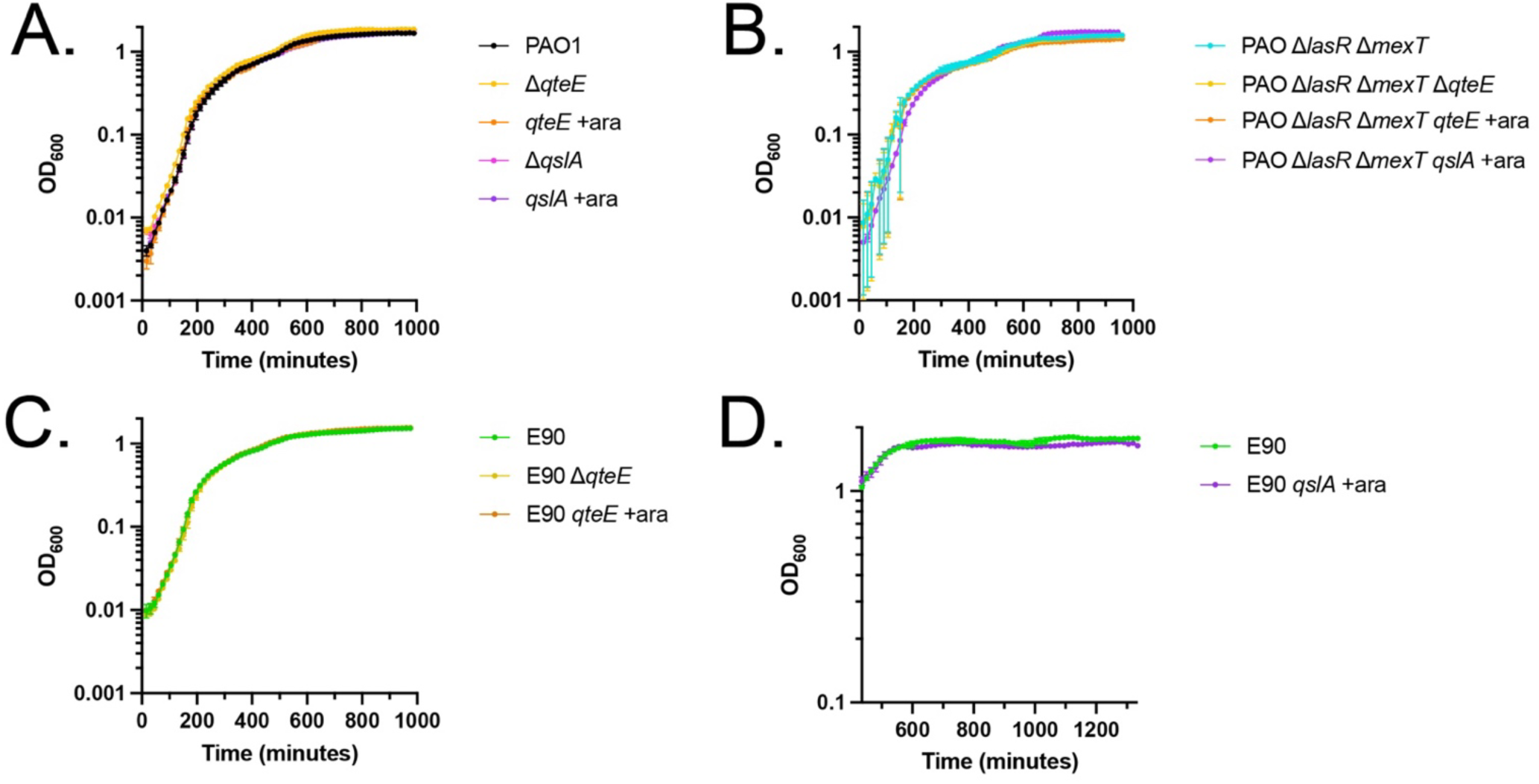
Growth curves of PAO1, PAO Δ*lasR* Δ*mexT*, and E90 as measured by OD_600_, under the experimental conditions shown in Figure 1. All data points show means of three biological replicates, and error bars represent s.e.m. Some error bars may not be visible.

**Figure S4:**
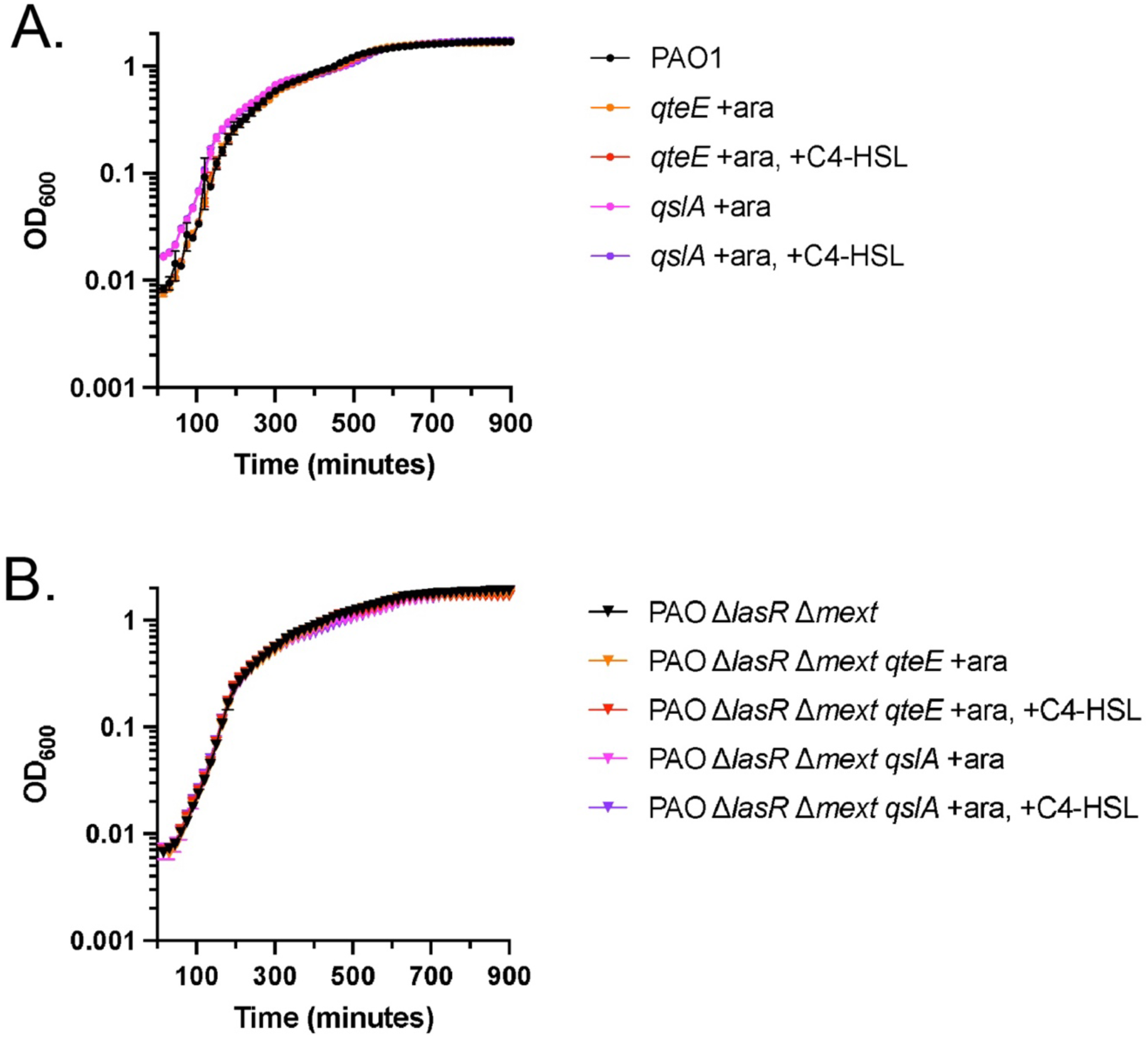
Growth curves of PAO1 (A) and PAO Δ*lasR* Δ*mexT* (B) strains tested in Figure 2, as measured by OD_600_. All data points show means of three biological replicates, and error bars represent s.e.m. Some error bars may not be visible.

